# DNA metabarcoding reveals limited consumption of livestock and black rhinoceros by spotted hyenas in a prey-rich environment

**DOI:** 10.1101/2023.10.19.563067

**Authors:** Arjun Dheer, Renita Danabalan, Antonia Pellizzone, Eve Davidian, Philemon Naman, Camila Mazzoni, Oliver P. Höner

## Abstract

The diet of large carnivores is of great interest to conservation managers, as it can reveal the extent of human-carnivore conflict and the impact of carnivores on species of high conservation priority. Metabarcoding of environmental DNA can identify species and is often more reliable in doing so than observational or morphological methods. Metabarcoding is particularly powerful at detecting elusive and rare species and has therefore become a widely applied tool in biodiversity research. Here, we used DNA metabarcoding of fecal samples to determine the diet of spotted hyenas in the Ngorongoro Crater, a protected area in Northern Tanzania surrounded by areas co-inhabited by pastoralists. We assessed which species hyenas preferably consumed over a 24-year period and how frequently they consumed pastoralist livestock and black rhinoceros, a species of high economic value and conservation priority. We further estimated the effects of three key socio-demographic variables – age, social rank, and sex – on the propensity of hyenas to consume livestock. We detected DNA from 20 species in 371 hyena feces. Hyenas preferably consumed blue wildebeest and Grant’s gazelle. Among the detected species were five domestic species and one wild species that lived in the pastoralist-inhabited areas but not the Crater. This shows that resident Crater hyenas undertake foraging trips to areas surrounding the Crater. DNA of domestic species however was rarely detected (4.1% of 434 detections), and predominantly in feces of old hyenas. This suggests that Crater hyenas rarely consume livestock and that livestock is mostly consumed by hyenas less capable of hunting fleet-footed and powerful wild prey. No DNA of black rhinoceros was detected in any of the samples, suggesting that Crater hyenas do not frequently consume rhinoceros. Our findings suggest that the impact of Crater hyenas on livestock and wildlife of high conservation priority is limited. Our study highlights the potential of DNA metabarcoding to assess the extent of human-carnivore conflict and to guide evidence-based conservation efforts to promote coexistence of carnivores, humans and species of high conservation priority.

## 1. Introduction

The effective management of multi-use protected areas (MUPA), where humans and wildlife cohabit, depends on gathering rigorous evidence to answer pressing conservation questions (Watson et al., 2016). Finding answers to such questions is especially urgent for the conservation of threatened wildlife species (Western, Russell, & Cuthill, 2009). Using such an evidence-based approach helps to balance the needs of priority wildlife species with those of local stakeholders, and the promotion of human-wildlife coexistence (Wevers et al., 2020).

There is currently a lack of evidence and conservation research conducted in protected areas, and particularly MUPA (Coetzee, 2017). This has hamstrung successful conservation and led to harmful management decisions, such as the broadly applied indiscriminate culling of animals that help neither people nor wildlife. For example, poorly justified culling of rodents (*Rodentia* spp.) may have weakened multiple ecosystem services, reduced the food base for predatory birds and mammals, and caused habitat loss for other burrowing animals (Singleton et al., 2007). Management decisions may also overlook individual-level tendencies in a population. For example, ‘problem animals’ may develop a predilection for consuming domestic animals, such as livestock. This behavior may be influenced by individual socio-demographic traits such as age, social rank, or sex (e.g., in brown bears (*Ursus arctos*): Morehouse et al., 2016). For instance, older age has been suggested to be positively associated with the killing of livestock or even humans by large carnivores (Reza, Feeroz, & Islam, 2002). This is because old individuals may be less capable of successfully hunting fleet-footed, powerful wild prey due to age-related disease and injury, and loss of dexterity, endurance, speed, and strength (Peterhans & Gnoske, 2001). In such cases, local communities may participate in indiscriminate retaliatory killing, or authorities may apply very broad conflict mitigation measures. While these actions may help reduce the perceived conflict in the short term, they can have major negative effects on populations and ecological communities, exacerbating the initial problem (Swan et al., 2017). There is accordingly an urgent need for using rigorous methods to gather robust evidence to answer important conservation questions.

DNA metabarcoding is a recently developed, non-invasive technique to identify species in multi-species samples, including environmental DNA (eDNA) samples. It has increasingly been used to assess biodiversity and determine the diet of wild carnivores (Berry et al., 2017; Havmøller et al., 2019; Morin et al., 2016; Ruppert et al., 2019). In carnivore research, scientists collect fecal samples from the carnivore and extract and identify the DNA of consumed species using high-throughput sequencing of taxonomically-informative markers (Alberdi et al., 2019). DNA metabarcoding poses several advantages over traditional diet assessments based on direct observation of consumed prey or morphological analyses of prey hair in feces. It often detects consumed taxa otherwise missed (Berry et al., 2017). It reduces representation bias associated with major differences in surface-area-to-volume ratios and hair densities across different species (Wachter et al., 2012) and can verify the identity of the putative consumer (Shehzad et al., 2015). It is also less likely to lead to misidentification between closely related domestic and wild taxa than morphological approaches do (Monterroso et al., 2018). Although DNA metabarcoding cannot distinguish between hunting and scavenging (Toju & Baba, 2018), distinguishing the two behaviors was not the focus of our study.

In this study, we used DNA metabarcoding of fecal samples collected over a period of 24 years to assess the diet of spotted hyenas (*Crocuta crocuta;* henceforth ‘hyena’) in the Ngorongoro Crater (henceforth ‘Crater’), a caldera within a large MUPA and UNESCO World Heritage Site. Hyenas are apex predators and scavengers ranging across much of sub-Saharan Africa (Bohm & Höner, 2015). The Crater hyena population has been the subject of a long-term study since 1996 (Höner, Davidian, & Szameitat, 2022). Past research has suggested that hyenas exhibit considerable age-, rank-, and sex-related behavioral differences (Benhaiem et al., 2012; Boydston et al., 2003; Davidian et al., 2021; Hofer & East, 1993; Höner et al., 2007, 2010; Yoshida et al., 2013), although this has not yet been assessed in terms of diet. We addressed three conservation questions of high importance to the local authorities and of broad interest to applied ecologists and wildlife managers. First, we determined which animal species hyenas preferably consumed and how frequently they left the Crater floor to consume pastoralist livestock in the areas surrounding the Crater. Second, we assessed how often they consumed black rhinoceros (hereafter ‘rhino’; *Diceros bicornis*), a critically endangered species and a species of great economic value for tourism (Emslie, 2020). Third, we investigated how socio-demographic variables affect the propensity of hyenas to consume domestic animals.

## 2. Methods

### 2.1 Study area

The Ngorongoro Crater (Figure 1) is located in the Ngorongoro Conservation Area (NCA), Tanzania (03°12′36″S 35°27′36″E), and is part of the greater Serengeti ecosystem. The NCA is a MUPA established in 1959 and a United Nations Educational, Scientific, and Cultural Organization (UNESCO) World Heritage Site with a double mandate to protect the interests of wildlife and local human communities (Charnley, 2005). The NCA is inhabited by the Maasai tribe, a semi-nomadic, pastoralist ethnic group ranging from central Kenya to southern Tanzania (Fratkin, 2001). The Maasai resided in the Crater until 1974, when they were evicted and required to live in other parts of the NCA (Moehlman et al., 2020). They were allowed to enter the Crater with livestock to conduct pastoralist activities during daytime until the end of 2016 (Melubo & Lovelock, 2019). All lab work occurred at the Leibniz Institute for Zoo and Wildlife Research and the Berlin Center for Genomics in Biodiversity Research, both located in Berlin, Germany.

**Figure 1:**
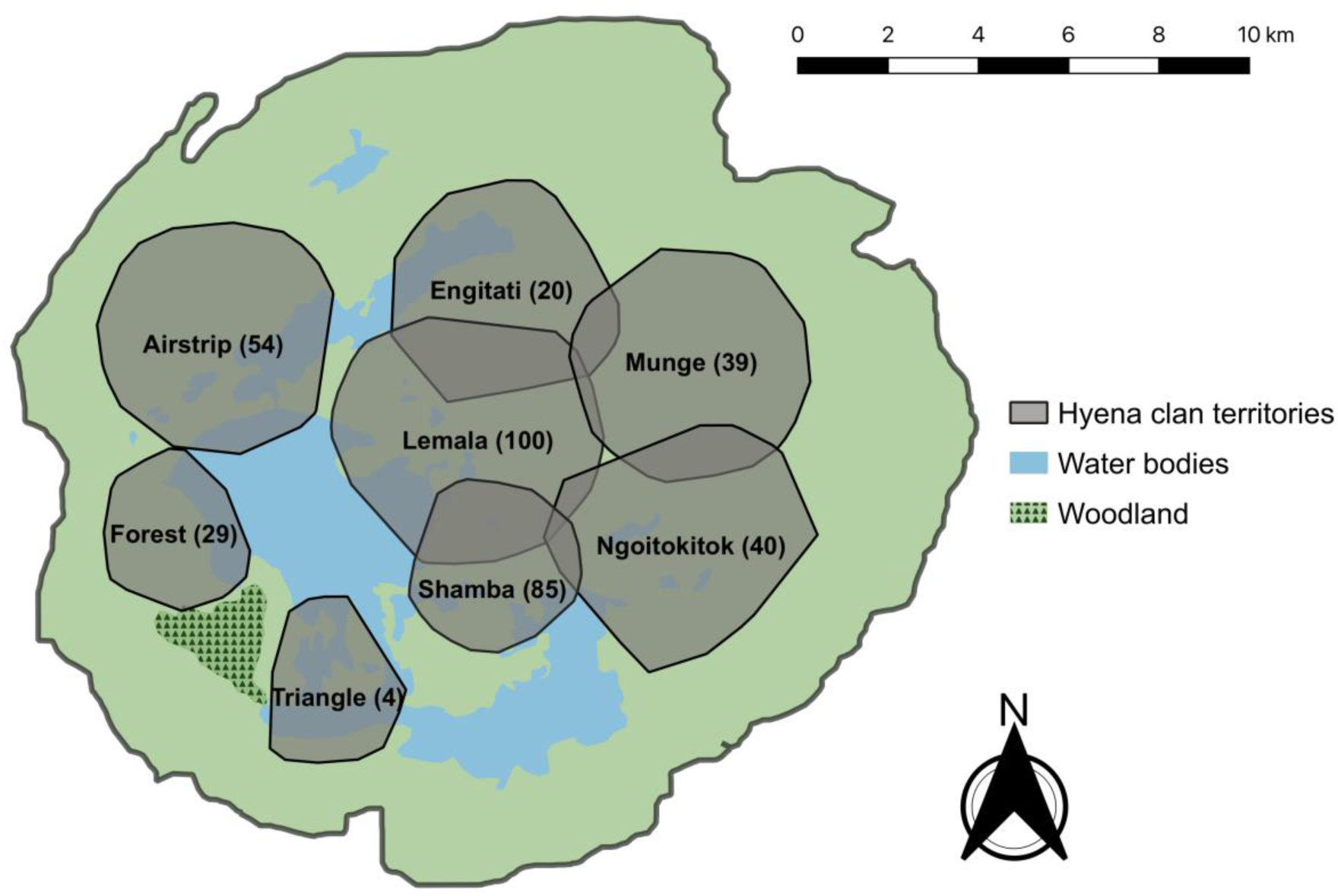
Location of the territories of the eight spotted hyena study clans in the Ngorongoro Crater. Territory boundaries are based on 85% minimum convex polygons (MCP) of adult female hyena sightings from 1996-2019 for each clan. MCP of 85% were chosen to accurately represent the locations and sizes of clan territories across the study period. Territories are labeled with corresponding clan names. Numbers in parentheses indicate the number of samples we analyzed from the corresponding clan.

Throughout the 24-year study period, the Crater was exceptionally dense in wild ungulates (Moehlman et al., 2020). In the areas surrounding the Crater, the populations of humans and their livestock greatly increased during this period. The human population increased from under 43,000 to over 100,000 (Manzano & Yamat, 2018), the cattle (*Bos taurus*) population more than doubled from under 120,000 to over 240,000, while the donkey (*Equus asinus*), goat (*Capra hircus*), and sheep (*Ovis aries*) populations increased to over 20,000, 220,000, and 340,000, respectively (National Bureau of Statistics Tanzania, 2017). Other domestic animals in the NCA included cats (*Felis catus*), chickens (*Gallus gallus*), dogs (*Canis familiaris*), dromedary camels (*Camelus dromedarius*), and pigs (*Sus domesticus*).

### 2.2 Data collection

Collection of field data occurred between April 1996 and December 2019, on a near-daily basis, between 06:00 and 19:00. We recognized individual hyenas based on their pelage patterns, ear notches, scars, and other traits. We estimated the age of cubs based on pelage, body size, behavior, and locomotion, with an accuracy of ±7 days (Pournelle, 1965) and identified the sex of individuals through observation of external genitalia (Frank, Glickman, & Powch, 1990). We collected fecal samples opportunistically, immediately after defecation by identified individuals. Following collection, we mechanically mixed and subsampled the feces, stored them in liquid nitrogen, and transported them to the laboratory in Berlin, Germany, on dry ice, where we stored them at −80°C until processing them for extraction and analyses. We conducted lab work between May 2019 and March 2021.

### 2.3 Molecular methods and socio-demographic representation

#### 2.3.1 DNA extraction

We extracted DNA from 505 fecal samples using a Stool DNA Kit (Roboklon GmBH, Berlin, Germany), following manufacturer instructions, with the following adjustments. We placed at least 200mg of feces into individual bead tubes from the kit and then homogenized the sample (Tissue Lyser II, Qiagen, Germany) two times for 20 seconds. We used the entire lysate for the remaining extraction procedure and eluted the samples in a final volume of 70µl. The concentration of extracted DNA was measured using a Qubit 3.0 Fluorometer (Thermo Fisher Scientific, Germany). We then stored samples in a −20°C freezer until sequencing took place.

#### 2.3.2 Sequencing summary

Out of the 505 fecal samples, we retained results from 371 (73.5%) for our analyses. Of the 134 samples we removed, 41 had low total read counts (<1,000 counts; Johnson, Lewandowski, & Merkes, 2021). Another 70 samples had low read counts and a DNA concentration that was too low (n = 63; <50 ng/µl) or too high (n = 7; >600 ng/µl) to be reliably sequenced (Koetsier & Cantor, 2019), possibly due to inhibitors preventing amplification of the target fragment (De Barba et al., 2013). The remaining samples that we removed (n = 23) were either duplicates derived from the same stool (i.e., samples collected from the same ‘defecation event’), or definite cases of contamination. For example, we had one sample with a total read count of ≥1,000, and most of its reads were of red fox (*Vulpes vulpes*), which are not extant in the NCA, let alone sub-Saharan Africa (Hoffman & Sillero-Zubiri, 2016) but samples of which were analysed in the same lab during the same period. Of the 371 samples we retained, 64 samples had read counts between 1,000-2,000, 95 samples between 2,000-3,000, and 212 samples >3,000.

#### 2.3.3 PCR amplification and sequencing

Because DNA in fecal samples can be heavily degraded, we used a 108-base pair (bp) fragment within the 12S rRNA gene (Riaz et al., 2011) with a two-step metabarcoding protocol. Briefly, vertebrate DNA was first amplified with primers containing Illumina adapters and a blocking primer specifically designed to inhibit the amplification of host DNA (i.e., hyena DNA; 5’-AGATACCCCACTATGCCTAGCCCTAAACTCAGATAGATAATT-Spacer_C3-3’). Depending on the DNA concentration, we used either 5µl or 200ng of DNA in a 20 µl PCR containing 1X Herculase buffer, 0.2nM of both dNTPs and forward and reverse primer, 10nM of blocking primer, 4nM of Mg2+, and 1.25U of Herculase II polymerase. Thermocycling conditions included an initial denaturation at 95°C for 5 minutes, followed by 35 cycles of 94°C for 30 seconds, 60°C for 30 seconds and extension at 72°C for 1 minute and ended with a 10-minute final elongation step at 72°C. We cleaned the PCR products using CLEAN NGS magnetic beads (GCBiotech, Netherlands) in a 0.8:1 ratio of beads to product and washed them twice, using 80% EOTH. We re-suspended products in 25µl of 0.1X T.E. buffer, and then recovered 24µl for the second PCR step. In order to de-replicate the individual samples post-sequencing, we PCR-ligated each sample with a unique combination of P7 and P5 indices, containing the sequencing primer and part of the flow cell adapter. As such, we added 10 µl of cleaned PCR product to a 15 µl mixture containing 1X Herculase buffer, 0.1nM dNTPs, 4% DMSO, 0.21U of Herculase II Polymerase and 0.25nM of both P7 and P5 indices. Thermocycling conditions included an initial denaturation at 95°C for 5 minutes, followed by 12 cycles of 94°C for 30 seconds, 52°C for 30 seconds, extension at 72°C for 1 minute, and ended with a 10-minute final elongation step at 72°C. We cleaned the now-indexed samples using the method described after the first PCR. Quantification of cleaned libraries were measured using a Quant-iT^TM^ Picogreen^TM^ dsDNA High Sensitivity Assay Kit (Thermo Fisher Scientific, Germany). Based on the concentration, we pooled samples equi-mass with 15ng of DNA. We cleaned the pool twice to remove any remaining primer dimers or sequencing products that were less than 100 bp, using the steps outlined above. We recovered a total of 20µl following the final elution. We checked libraries for a single fragment size of 320 bp using an Agilent Tapestation (Applied Biosystems, Germany) and quantified in replicate using a Qubit 3.0 Fluorometer (Thermo Fisher Scientific, Germany). We carried out sequencing on a MiSeq, v3 600 Cycles Reagent Kit (Illumina, U.S.A.).

#### 2.3.4 Quality control and analysis of sequences

We assessed all reads for quality using ‘FASTQC’ (Babraham Bioinformatics, England) and ‘MultiQC’ software (Ewels et al., 2016). The number of reads generated per sample was determined from the R1.fastq.gz and samples with a total read count of ≥1,000 (Clarke et al., 2020) were retained. We assembled read 1 and read 2 and removed poor quality sequences and sequencing adapters using Paired End reAd mergeR (PEAR, HITS, Heidelberg, Germany), keeping only reads with quality ≥18, based on the MultiQC output, by defining it with parameter: *-q 18*. We then removed primers using cutadapt: *-g forwardprimer…reverseprimer*, specifying that adapters are linked and requiring both to be removed (Martin, 2011). We removed chimeras using abundance rather than a reference with *Uchime_denovo* with the ‘Vsearch’ programme (Rognes et al, 2016) after de-replication. Following the removal of chimeras, we re-replicated samples and appended sample names into the header of the sequence using *Obiannotate –sample S:sample_name*, within the ‘OBItools’ programme (Boyer et al., 2014). We de-replicated sequences again using *Obiuniq – m sample*, as the output contained the number of reads for each unique sequence cluster. Based on the read counts in the *Obiuniq* output, we only retained sequences with read counts of ≥5% of the total count within the sample (‘discardPercentageofmaxderep_Fasta.py’ script in Supplementary Material). Finally, we allocated species assignments using a BLASTn search on Genbank, which contains a reference library of over 450,000 formally described species (Sayers et al., 2020), including all terrestrial mammals known to be extant in the NCA (Moehlman et al., 2020). We assumed each individual detection of a given species’ DNA in a sample represented one ‘meal’, i.e., one case of the hyena eating an animal of that species (Davidson et al., 2019; McLennan et al., 2022).

#### 2.3.5 Verification of species identification

To validate our method to distinguish between the DNA of cattle, the most important livestock species to the Maasai community (Goldman, 2011), and the closely related African buffalo (*Syncerus caffer*; henceforth ‘buffalo’), we conducted a feeding experiment with a hyena housed at Tierpark Berlin, a zoo in Berlin, Germany, in 2019. On a designated morning, the hyena keeper fed the hyena exclusively cattle meat (beef). The next morning, based on the speed of the hyena digestive process (4-48 hours; authors’ observation; Goymann et al., 2019), the keeper collected a fecal sample from the hyena. In this sample, the extracted DNA was clearly identified as cattle DNA. In addition, we were able to verify the species identification three times based on direct field observations. First, ≥5% of reads of a fecal sample collected on 1998-07-07 from a hyena that ate blue wildebeest (*Connochaetes taurinus*; henceforth ‘wildebeest’) on 1998-07-06, were from wildebeest. Second, a sample from a hyena that we saw eating from a wildebeest carcass six hours prior to sample collection contained only wildebeest DNA. Third, the feces collected on 2012-11-29 from a hyena that we observed eating from a buffalo on 2012-11-28 contained buffalo and wildebeest DNA (≥5% of the reads for both species). We did not see this hyena eating wildebeest on 2012-11-28, but it is feasible that it did so at a time when we were not monitoring it (e.g., at night, given that we were restricted to daytime observations).

### 2.4 Age, social rank, sex, and consumed species category

The 371 analyzed samples were from 255 individual hyenas that were ≥1 year old on the date of collection. The hyenas represented all eight Crater clans (Figure 1). 234 samples (63.1%) were from males and the remaining 137 from females. The collection year of samples ranged from 1996 (n = 9) to 2019 (n = 39), with a peak in 2018 (n = 78; Figure S1). We attributed to each sample a unique ID as well as the sex, age and social rank of the corresponding defecating hyena on the day of sample collection. We determined ordinal ranks based on the history of recorded agonistic interactions and our knowledge of rank inheritance and social queuing (for details, see Davidian et al., 2021). We converted the ordinal rank (*OrdRanki*) of an individual (*i*) into a proportional rank (*PropRanki*) bounded between −1 (bottom rank) and 1 (top rank), accounting for clan size *N*, using the following formula:

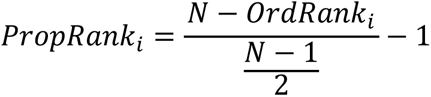

We estimated the effects of hyena age (in years: numeric and continuous), social rank (proportional: between −1 and 1), and sex (categorical: female or male) on the propensity of hyenas to consume domestic animals using a generalized linear mixed-effects model (GLMM) with a binomial distribution using package ‘lme4’ (Bates et al., 2015). We used the binomial distribution because the response variable was categorical, with two levels: ‘domestic’ or ‘wild’, depending on whether the given detection was of a domestic or wild animal species. We included a random effect for the ID of the fecal sample, given that some samples had more than one species detected.

We computed predictions and associated 95% confidence intervals from the GLMM for any significant independent variable(s) using the function ‘ggemmeans’ from package ‘ggeffects’ (Lüdecke, 2018). The coefficients estimated by the model were log(odds) and were converted into odd ratios, using the formula: exp[coefficient]. Odd ratios > 1 and odd ratios < 1 indicate a relative increase and decrease, respectively, in the likelihood to express the behavior.

### 2.5 Feeding preferences

We used data from biannual, transect-based Crater censuses (n = 43; Moehlman et al., 2020) to estimate the composition and size of the typically available food base for the hyena population over the course of our study period. We determined feeding preferences for the different species by calculating Manly’s standardized selection ratio *B* (Manly, McDonald, & Thomas, 1993) using package ‘adehabitatHS’ (Calenge, 2006). To ensure our estimates of *B* were rigorous (Manly, McDonald, & Thomas, 1993), we restricted analyses to the five species for which there were reliable census data and at least 10 detections; wildebeest, buffalo, zebra (*Equus quagga*; henceforth ‘zebra’), Grant’s gazelle (*Nanger granti*), and Thomson’s gazelle (*Eudorcas thomsonii*), the five most commonly hunted herbivores by Crater hyenas (Höner et al., 2002). The ratio *B* provides an estimate of selection for a given resource (i.e., the consumed species) relative to selection for all other resources, based on the proportional availability of each resource within the entire community. The value of *B* is the probability that a randomly chosen detection of any species will belong to a given species, if all species are equally frequent in the original population (Manly, McDonald, & Thomas, 1993). Thus, the ratio can range from 0 (minimum selection) to 1 (maximum selection). To determine if the proportional consumption of each species was significantly different from its corresponding proportional availability, we used a chi-square test with Bonferroni-corrected p-values.

### 2.6 Statistical analyses

We conducted all statistical analyses in R software 4.2.0 (R Core Team, 2022). Data are presented as means ± S.D., unless stated otherwise. The threshold for statistical significance was set to *α* = 0.05.

## 3. Results

### 3.1 Animal species detected in spotted hyena feces

We detected 20 different animal species in 371 hyena feces (Figure 2, Table S1). The total number of detections – inclusive of all species – was 434. The majority of feces (n = 312; 84.1%) only contained one species, the others either contained two (n = 55) or three species (n = 4) (Table S1).

**Figure 2:**
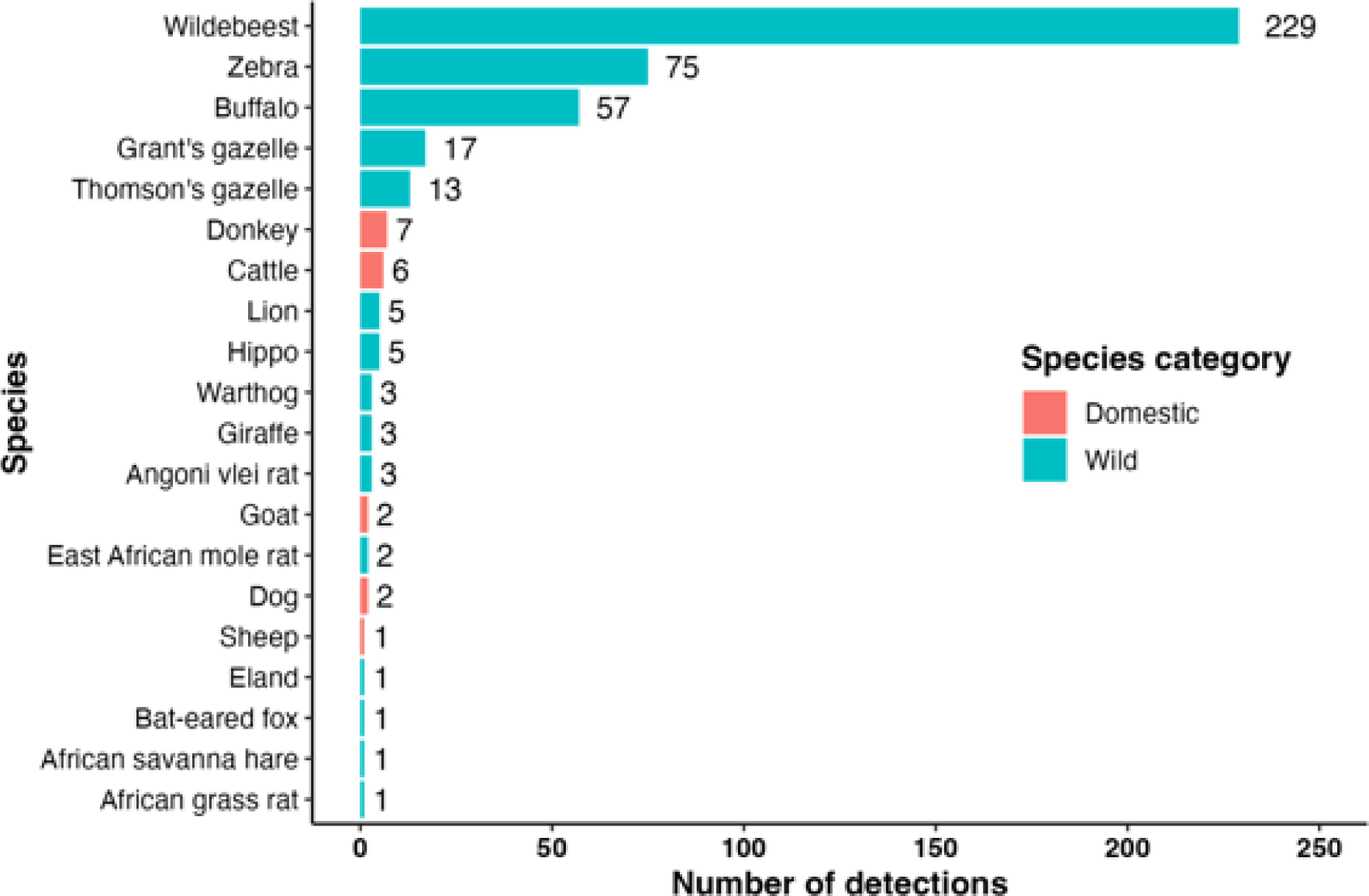
Total number of detections of DNA of different animal species in spotted hyena feces from the Ngorongoro Crater. There were 434 detections across 371 feces. Each detection represents one case of DNA of the given species being detected. Detections were classified as ‘domestic’ (domestic animal species) or ‘wild’ (wild animal species) and are color-coded accordingly.

Wildebeest was by far the most commonly detected species, followed by zebra, and buffalo. There were no detections of rhino and three detections of Maasai giraffe (*Giraffa camelopardalis tippelskirchi*; henceforth ‘giraffe’), a species only extant outside the Crater. Domestic animals (cattle, dog, donkey, goat, and sheep) were detected 18 times, which corresponds to 4.1% of all detections.

### 3.2 Detections according to socio-demographic variables and species categories

The hyenas in whose feces domestic animal DNA was detected were older (mean age = 9.08 ± 4.16 years, n = 18 detections) than the hyenas that consumed wildlife species (6.19 ± 3.04 years, n = 416; two-sample permutation test; Z = 3.82, p < 0.001; Figure 3A). The proportional rank of hyenas consuming domestic (−0.39 ± 0.62) and wild (−0.23 ± 0.63) animals was similar (Z = −1.06, p = 0.29; Figure 3B).

**Figure 3:**
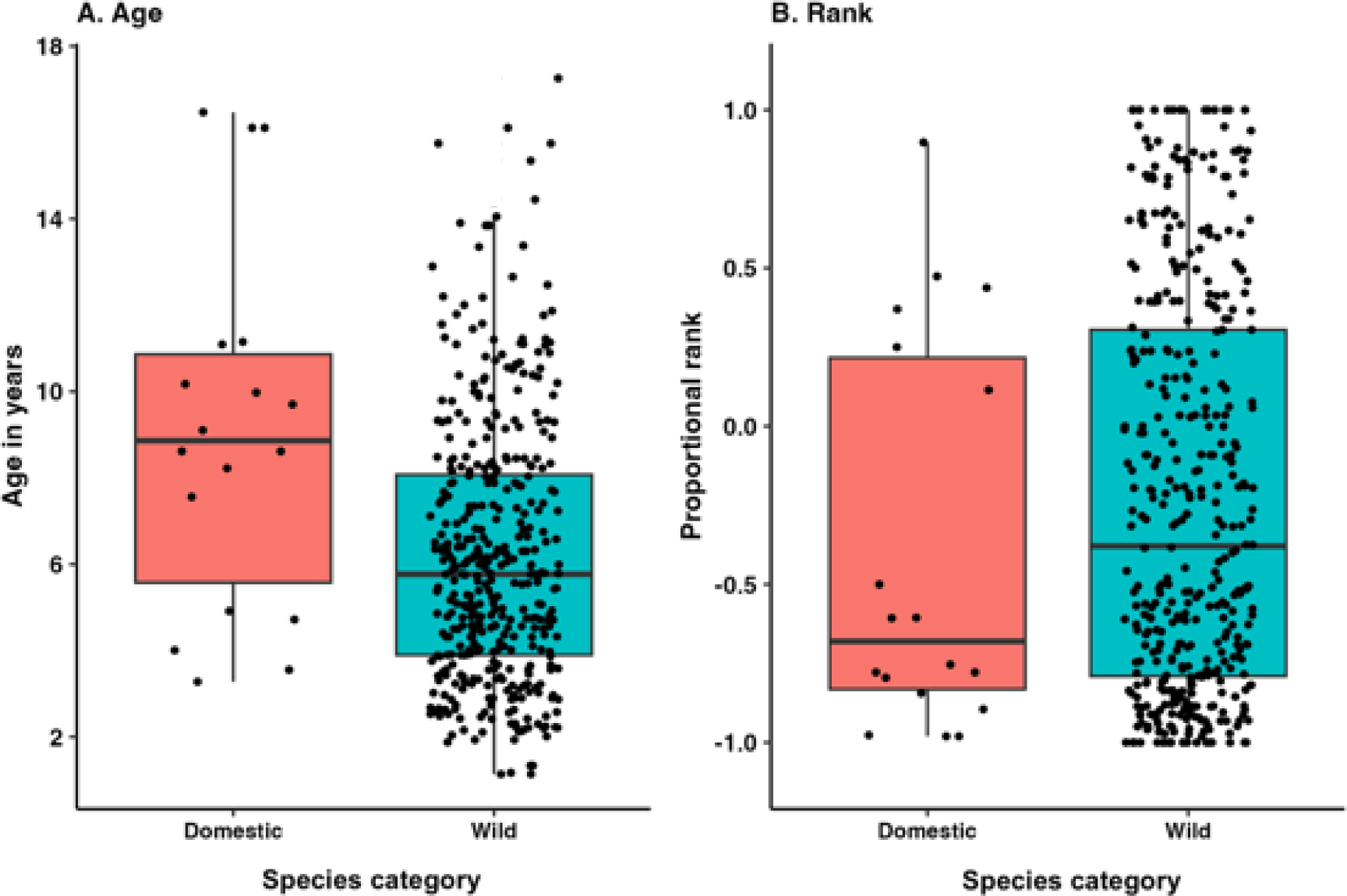
Consumption of domestic and wild animals by spotted hyenas from the Ngorongoro Crater as a function of hyena age (A) and social rank (B). Boxes indicate the interquartile range around the median (horizontal bar), vertical bars represent data that lie within 1.5 times the interquartile range. Black dots represent observed data after horizontal jittering was applied for ease of visualization.

The majority of detections (n = 273; 62.9% of all detections) were from males. Similarly, the majority of both domestic (n = 13; 72.2%) and wild (n = 260; 62.5%) animal detections also came from males.

#### 3.2.1 Model results

There was a significant positive effect of age, but not of social rank or sex, on the predicted probability of domestic animal consumption (Table 1). A one-year increase in a hyena’s age led to a predicted 26.8% increase in the probability of domestic animal consumption.

**Table 1:** Effects of socio-demographic variables on the propensity of spotted hyenas from Ngorongoro Crater to consume domestic animals. Covariates consist of age, social rank, and sex. The intercept for the model corresponds to the female sex, with age and rank held at their means. The column “S.E.” provides standard errors on parameter estimates. The column “Odd ratios” corresponds to the exp(estimate) and indicates the factor of change in the likelihood to consume domestic animals relative to that of consuming wild animals, with a one-unit increase in the predictor and when other covariates are held constant. The columns “Z” and “*p*” give, respectively, the Z- and p-values associated with the likelihood ratio test. Data in bold were deemed significant. Results are based on a binomial GLMM. Random variance was estimated for sample ID (n = 371 samples) at 0.74.

|  | **Estimate** | **S.E.** | **Odd ratios** | **Z** | **p** |
| --- | --- | --- | --- | --- | --- |
| Intercept | -5.35 | 0.87 |  |  |  |
| Age | **0.24** | **0.07** | **1.30** | **3.29** | **0.001** |
| Social Rank | -0.07 | 0.63 | 0.93 | -0.11 | 0.91 |
| Sex Male | 0.55 | 0.83 | 1.73 | 0.66 | 0.51 |

### 3.3 Dietary preferences

Wildebeest was selected for the most strongly, followed by Grant’s gazelle, buffalo, zebra, and then Thomson’s gazelle (Table 2).

**Table 2:** Dietary preferences by spotted hyenas in the Ngorongoro Crater. Abundances were estimated based on transect surveys conducted within designated census blocks from 1996-2019 by the Ngorongoro Conservation Area Authority, usually bi-annually. Presented are results for the five species for which abundance estimates existed and that were detected ≥10 times. Column “*B*” provides Manly’s standardized selection ratio for the given species, with higher values indicating greater selection for the given species. Columns “χ^2^” and “*p*” provide the chi-square test statistic and Bonferroni-corrected p-value, respectively, for the proportional consumption and availability of each species. Column “Sign” shows whether the proportional consumption was significantly greater than expected (+), less than expected (−), or as expected (0). Data in bold were deemed statistically significant.

| Species | Abundance ± S.D. | *B* | $\chi^2$ | *p* | Sign |
| --- | --- | --- | --- | --- | --- |
| Wildebeest | **8,218.8 ± 2,924.4** | **0.30** | **25.75** | **<0.001** | + |
| Grant's gazelle | **833.7 ± 396.4** | **0.22** | **5.58** | **0.018** | + |
| Buffalo | 3,176.8 ± 1,540.1 | 0.19 | 3.16 | 0.075 | 0 |
| Zebra | 4,274.1 ± 1,331.1 | 0.19 | 0.09 | 0.76 | 0 |
| Thomson's gazelle | **1,392.4 ± 632.0** | **0.10** | **24.15** | **<0.001** | - |

## 4. Discussion

In this study, we used DNA metabarcoding to assess the diet of spotted hyenas in the Ngorongoro Crater over a 24-year period. We sought to estimate the level of conflict between hyenas and humans by determining how often Crater hyenas consumed pastoralist livestock and wildlife of high economic value and conservation priority. We further aimed at identifying traits that predispose hyenas to come into conflict with humans. Overall, the diet of Crater hyenas consisted mainly of wild animals, with only few detections of domestic animal DNA in hyena feces. The likelihood of hyenas to consume domestic animals increased with increasing age. No DNA of black rhinoceros was detected in any of the hyena feces, suggesting that Crater hyenas do not frequently consume this critically endangered species. Below, we provide possible explanations for and interpretations of our results in the context of evidence-based large carnivore conservation.

The diet of Crater hyenas predominantly consisted of 15 species of wild animals, with wildebeest being the most strongly preferred species, followed by Grant’s gazelle. These results are similar to those of Höner et al. (2002), who analyzed direct observations of hunts and found that Crater hyenas preferably hunted wildebeest and juvenile gazelles (Grant’s and Thomson’s combined). One of the species detected in the current study was giraffe, a species of wild herbivore that lives in the areas surrounding the Crater but not on the Crater floor. This shows that resident Crater hyenas occasionally undertake foraging trips to areas outside the Crater, corroborating indirect evidence from a previous study (Höner et al., 2005).

The domestic animals whose DNA was detected in Crater hyena feces also most likely originated from outside the Crater because Maasai pastoralists and their livestock do not reside in the Crater. Livestock were permitted to enter the Crater during daytime until 2017, but they were very well protected by the pastoralists (Dheer et al., 2022). The small number of detections of livestock suggests that Crater hyenas rarely consume livestock, despite the high abundance of livestock in the areas surrounding the Crater. Furthermore, the fact that the detections include both depredation and scavenging of dead livestock and that there were frequent outbreaks of fatal diseases among livestock during the study period (Dheer et al., 2021) suggests that livestock depredation by Crater hyenas overall is very rare. It seems likely that the high abundance of wild herbivores in the Crater makes hunting and scavenging from wild animals relatively easy. This would be consistent with past research, which suggested that large carnivores only regularly consume domestic animals when wild prey is very scarce (Khorozyan et al., 2015; Parsons, Newsome, & Young, 2022; Yirga et al., 2012). Our results have implications for conservation management in the Crater and surrounding areas as they suggest that most depredation leading to human-hyena conflict in the NCA (Dheer et al., 2021) is by hyenas from outside the Crater and not by hyenas resident in the Crater.

Interestingly, we found a positive effect of age on the propensity of Crater hyenas to consume domestic animals. This may be because older hyenas are less likely to successfully hunt wild prey and more likely to know where to find and scavenge or hunt livestock. Past reports of culled ‘man-eaters’ or ‘problem animals’ suggested that old and injured large carnivores may be forced to rely on scavenging and feeding on domestic animals and humans due to age- and injury-related physical impairments (Rabinowitz, 1986). Other studies suggested that conflict-prone individuals (at least in large felids; *Felidae* spp.) tend to be young transients who have not yet established territories with sufficient wild prey (Patterson et al., 2003; Woodroffe & Frank, 2005). This is less likely in spotted hyenas, as females are usually resident in their natal clans for life (Höner et al., 2007), most males disperse only as adults (at around 3.5 years of age; Höner et al., 2010; Davidian et al., 2016), and transience generally is rare. In the Crater, only 15 of 470 males who joined a Crater clan were transient at some point in their life, and transient individuals were on average 7.9 ± 3.9 years old at the beginning of the transience period. These findings warrant further research into the effects of individual socio-demographic traits on the consumption of domestic animals by large carnivores. This can inform evidence-based large carnivore conservation actions tailored towards preventing specific individuals or classes of individuals from entering areas where they are likely to come into conflict with humans (Snijders et al., 2019).

The absence of detections of rhino DNA in hyena feces suggests that the Crater hyenas did not regularly consume rhinos. The potential effect of hyena predation on rhinos has been raised as a major conservation issue in the NCA and more globally in East Africa, but such concerns have lacked robust evidence (Davidson et al., 2019; Sillero-Zubiri & Gotelli, 1991). Our results may further assuage such concerns. While hyenas are known to scavenge and occasionally even attack and kill rhinos (Kruuk, 1972; Owen-Smith & Mills, 2008), the lack of rhino DNA in 371 hyena feces indicates that it is very unlikely to occur regularly. Our results therefore suggest that hyenas do not pose a credible threat to the persistence of the black rhino population in the Ngorongoro Crater. This is supported by a recent study that showed that the Crater rhino population increased substantially and more strongly during our study period than predicted under favorable conditions (Moehlman et al., 2020). We consider this a positive sign of the rhino population’s continued persistence, and the result of ongoing conservation efforts geared towards the protection of rhinos by the NCAA, including intensive surveillance and anti-poaching operations, and an evident lack of significant predation pressure by hyenas and lions (*Panthera leo*), the other main carnivore in the Crater.

Our study highlights the potential of DNA metabarcoding to assess the extent of human-carnivore conflict and to guide evidence-based conservation efforts to promote coexistence of carnivores, humans and species of high conservation priority.

## Acknowledgments

We thank the Tanzania Commission for Science and Technology, the Tanzania Wildlife Research Institute, and the Ngorongoro Conservation Area Authority for permission to conduct the study. Additionally, we thank B. Wachter for collecting behavioral and demographic data and fecal samples, M. Quetstroey for extracting DNA, and M. Driller for writing the ‘discardPercentageofmaxderep_Fasta.py’ script. We also thank L. Bailey, A. Courtiol, E. Donati, and Z. Li for co-developing the *hyenaR* package which was essential for data extraction. Finally, we thank the Leibniz Institute for Zoo and Wildlife Research and our supporters on Experiment.com for providing financial support.

