## Supplementary Material for "DNA metabarcoding reveals limited consumption of livestock and black rhinoceros by spotted hyenas in a prey-rich environment"

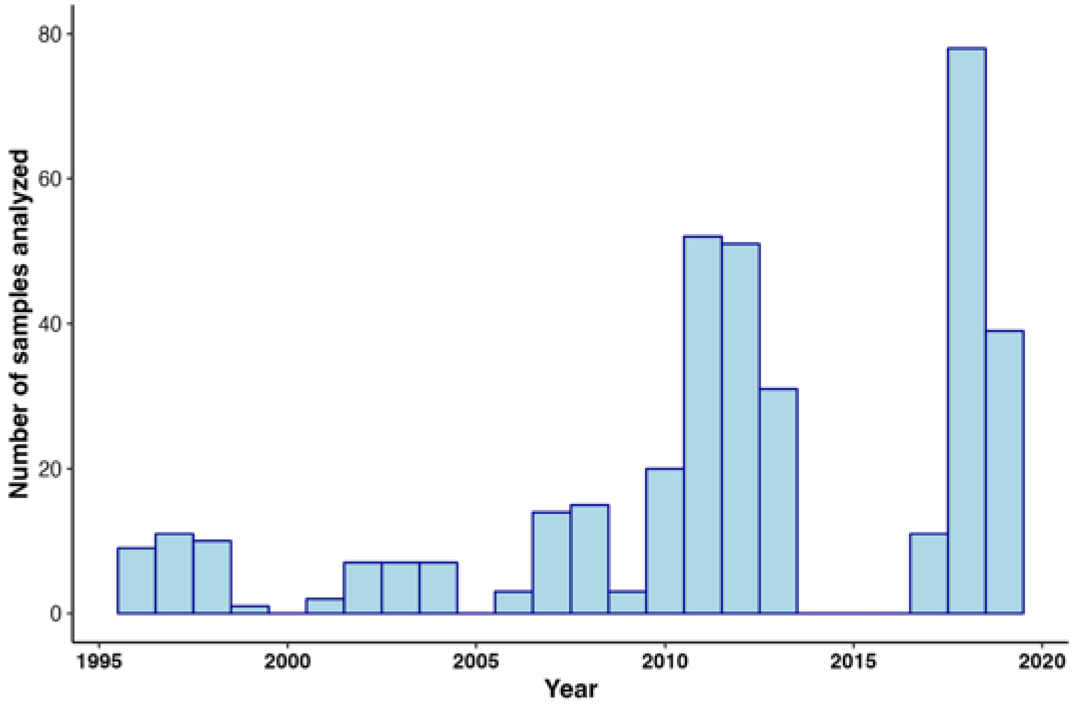


**Figure S1: Histogram displaying yearly distribution of fecal samples of spotted hyenas from the Ngorongoro Crater included in this study**. A total of 371 samples were used.

**discardPercentageofmaxderep_Fasta.py** script is provided below.

```
import argparse

parser = argparse.ArgumentParser(description="Parses a fasta file dereplicated by obitools and discard reads with less than a given %% of the maximum abundance (5%% default)")
parser.add_argument("-fa", "--fasta", help="fasta file of dereplicated reads using obitools")
parser.add_argument("-oa", "--outaccepted", help="output for accepted reads")
parser.add_argument("-od", "--outdiscarded", help="output for discarded reads")
parser.add_argument('-p', "--percentage", type=float, default=5.0, help="Percentage of total abundance to discard [default: 5%%]")
parser.add_argument("-max", "--maxAbundance", action="store_true", help="enable if using the max abundance instead of total to calculate min abundance to keep a sequence")
args = parser.parse_args()


faParser=open(args.fasta)


########## getting max abundance and calculating 5%
maxAbundance=0
totalAbundance=0
for line in faParser:
    if line.startswith(">"): # new read
        seq=""
        #print(line)
        readCount=int(line.split("count=")[1].split(";")[0]) # get the count value for read
        #print(readCount)
        totalAbundance+=readCount
        if readCount>maxAbundance:
            maxAbundance=readCount


faParser.close()

if args.maxAbundance:
    minAbundance=float(maxAbundance)/100*args.percentage
    print("Maximum abundance is: %i | Stacks with less than %f(%f%%) counts will be discarded"%(maxAbundance, minAbundance, args.percentage))
else:
    minAbundance=float(totalAbundance)/100*args.percentage
    print("Total abundance is: %i | Stacks with less than %f(%f%%) counts will be discarded"%(totalAbundance, minAbundance, args.percentage))


######## filtering based on max abundance
aWriter=open(args.outaccepted, "w")
dWriter=open(args.outdiscarded, "w")
```

```
#counters
seqCount=0
seqAcc=0
seqDis=0

readHeader="" #init
readSeq="" #init
readCount=0 #init
faParser=open(args.fasta)
write=False
for line in faParser:
    if line.startswith(">"): # new read
        write=False
        readCount=int(line.split("count=")[1].split(";")[0]) # get the count value for read
        if readCount>=minAbundance:
            aWriter.write(line)
            seqAcc+=1
            write=True
        else:
            dWriter.write(line)
            seqDis+=1

        seqCount+=1
        readSeq=""
        #print(line)
        readHeader=line
        readCount=int(line.split("count=")[1].split(";")[0]) # get the count value for read
        #print(readCount)
        if readCount>maxAbundance:
            maxAbundance=readCount
    else:
        if write==True:
            aWriter.write(line)
        else:
            dWriter.write(line)


faParser.close()
aWriter.close()
dWriter.close()


print("Of %i sequences in total %i were accepted and %i were removed"%(seqCount, seqAcc, seqDis))
```
